# Simplifying MS1 and MS2 spectra to achieve lower mass error, more dynamic range, and higher peptide identification confidence on the Bruker timsTOF Pro

**DOI:** 10.1101/2021.10.18.464737

**Authors:** Daryl Wilding-McBride, Laura F. Dagley, Sukhdeep K Spall, Giuseppe Infusini, Andrew I. Webb

**Affiliations:** The Walter and Eliza Hall Institute of Medical Research, 1G Royal Parade, Parkville, Melbourne, Victoria 3052, Australia; Department of Medical Biology, University of Melbourne, Parkville, Melbourne, Victoria 3010, Australia; Mass Dynamics, Melbourne, Victoria, Australia

**Author notes:** Corresponding authors Corresponding author details: Daryl Wilding-McBride, Andrew Webb.

## Abstract

1

For bottom-up proteomic analysis, the goal of analytical pipelines that process the raw output of mass spectrometers is to detect, characterise, identify, and quantify peptides. The initial steps of detecting and characterising features in raw data must overcome some considerable challenges. The data presents as a sparse array, sometimes containing billions of intensity readings over time. These points represent both signal and chemical or electrical noise. Depending on the biological sample’s complexity, tens to hundreds of thousands of peptides may be present in this vast data landscape. For ion mobility-based LC-MS analysis, each peptide is comprised of a grouping of hundreds of single intensity readings in three dimensions: mass-over-charge (m/z), mobility, and retention time. There is no inherent information about any associations between individual points; whether they represent a peptide or noise must be inferred from their structure. Peptides each have multiple isotopes, different charge states, and a dynamic range of intensity of over six orders of magnitude. Due to the high complexity of most biological samples, peptides often overlap in time and mobility, making it very difficult to tease apart isotopic peaks, to apportion the intensity of each and the contribution of each isotope to the determination of the peptide’s monoisotopic mass, which is critical for the peptide’s identification.

Here we describe four algorithms for the Bruker timsTOF Pro that each play an important role in finding peptide features and determining their characteristics. These algorithms focus on separate characteristics that determine how candidate features are detected in the raw data. The first two algorithms deal with the complexity of the raw data, rapidly clustering raw data into spectra that allows isotopic peaks to be resolved. The third algorithm compensates for saturation of the instrument’s detector thereby recovering lost dynamic range, and lastly, the fourth algorithm increases confidence of peptide identifications by simplification of the fragment spectra. These algorithms are effective in processing raw data to detect features and extracting the attributes required for peptide identification, and make an important contribution to an analytical pipeline by detecting features that are higher quality and better segmented from other peptides in close proximity. The software has been developed in Python using Numpy and Pandas and made freely available with an open-source MIT license to facilitate experimentation and further improvement (DOI 10.5281/zenodo.6513126). Data are available via ProteomeXchange with identifier PXD030706.

**Author Summary:** The primary goal of mass spectrometry data processing pipelines in the proteomic analysis of complex biological samples is to identify peptides accurately and comprehensively with abundance across a broad dynamic range. It has been reported that detection of low-abundance peptides for early-disease biomarkers in complex fluids is limited by the sensitivity of biomarker discovery platforms (1), the dynamic range of plasma abundance, which can exceed ten orders of magnitude (2), and the fact that lower abundance proteins provide the most insight in disease processes (3). As mass spectrometry hardware improves, the corresponding increase in amounts of data for analysis pushes legacy software analysis methods out of their designed specification. Additionally, experimentation with new algorithms to analyse raw data produced by instruments such as the Bruker timsTOF Pro has been hampered by the paucity of modular, open-source software pipelines written in languages accessible by the large community of data scientists. Here we present several algorithms for simplifying MS1 and MS2 spectra that are written in Python. We show that these algorithms are effective to help improve the quality and accuracy of peptide identifications.

## 3 Introduction

The task of a feature detector in an analytical pipeline is to sift through raw points and find the characteristic pattern of a peptide. Once a peptide feature is detected, the attributes that are important for its identification must be determined.

Most commonly, feature detection of mass spectrometry data has been designed to utilise three dimensions: intensity (the unit-less intensity dimension being a proxy for abundance), mass-over-charge (m/z) and retention time (4). The timsTOF mass spectrometer (5) adds the dimension of collisional cross section (CCS). This extra dimension gives an opportunity to separate peptides that are isobaric but have different collisional cross-sectional area (CCS). Peptide features in timsTOF raw data are thus represented in four dimensions: m/z, retention time, intensity, and CCS. While the added mobility dimension adds considerable information about the peptides’ unique characteristics, it results in much more data to process; a typical sample analysis using a 20-minute LC gradient generates 1.3 billion MS1 and 37 billion MS2 raw intensity readings.

Commonly used free-to-download tools for processing timsTOF data include MaxQuant (6), MSFragger (7,8), Biosaur (9), and recently AlphaPept (10). Biosaur and AlphaPept are in addition open-source software. These tools have established ecosystems around them and, to varying degrees, offer a means by which modules can be developed to interface with and extend them. As MaxQuant and MSFragger are closed source, it is not possible to propose code adjustments to improve performance, and contributions of feature extensions to the core modules are not possible. To understand the implementation details of Biosaur and AlphaPept, the source code is available to study, but for MaxQuant and MSFragger a description of the algorithm in prose form from the paper must be relied upon, which may be open to ambiguity or misunderstanding of important details.

In setting out to develop ideas for new acquisition modes and experimenting with new ideas that leveraged the timsTOF’s sensitivity, we perceived a shortage of raw data processing tools that had the following characteristics:

1. Open-source code published on a modern code sharing platform such as GitHub, BitBucket, or GitLab, so anyone with coding skills could inspect the details of the algorithm implementation, to understand the inner workings and perhaps to offer improvements to the implementation.
2. Open and well-documented inter-module interfaces designed for programmatic consumption and ingestion, taking modern payload formats such as JSON, to facilitate a plug-and-play architecture in which add-ons could be developed for alternative data processing, or extensions to the processing pipeline.
3. Implemented in a popular, accessible language such as Python and distributed as packages that can be installed and version-managed with standard tools for the language environment, such as `conda` or `pip` in the case of Python.

Existing software tools that are freely downloadable but lack one or more of these characteristics contribute to the processing of data but forego the opportunity to help progress the understanding of data processing algorithms, which means:

- Writing add-ons for customised workflows involves spending valuable effort writing code for parsing or generating files that are not intended for programmatic interfaces. For example, MaxQuant’s table text files (11) are textual representations of results tables that are more suited to human consumption than programmatic consumption and cross-linking.
- The pace of improvement in the processing of data is limited by the capacity of the original authors to make improvements.

Our aim was to address these issues by developing an open source timsTOF data processing pipeline from scratch, offering a software foundation that facilitates collaboration in the conduct of further research, covering topics including signal processing, feature detection and segmentation, peptide identification, performance optimisation, peptide quantification, and protein inference.

## 4 Results

The high-level steps involved in identifying peptides from the raw instrument database are shown in Figure 1. The completeness of the peptide identifications produced depends on the precision of the preceding steps; from detecting features in the raw data, to determining the characteristics of those features, to extracting the fragment ions associated with the precursor ions and presenting those characteristics to a peptide database search.

**Figure 1.**
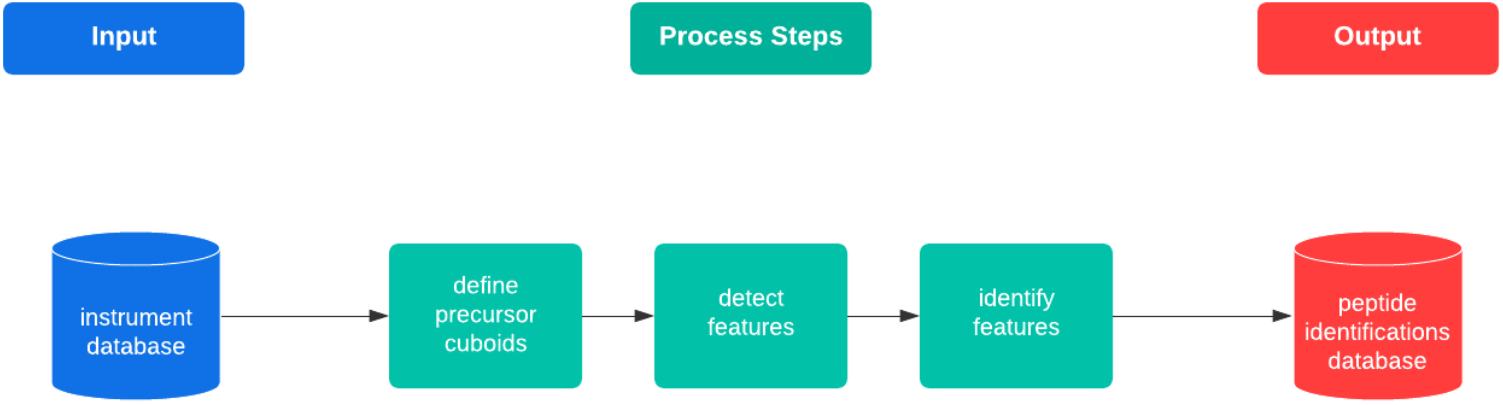
the high-level flow of the pipeline steps to process raw timsTOF data and identify peptide sequences

The following sections 4.1 to 4.4 describe four algorithms we developed to simplify the complexity of the MS1 and MS2 spectra derived from the raw data and to improve the quality of peptide identifications.

### 4.1 Finding peptide features in raw data

During acquisition, the instrument selects precursor ions for fragmentation based on their intensity. The windows selected for fragmentation in each frame are called isolation windows: small regions in the plane formed by the m/z and mobility dimensions. As a precursor ion elutes and increases in intensity, the instrument will select it for fragmentation multiple times. This results in multiple isolation windows across several frames. These isolation windows are used to schedule fragmentation events, where all ions in a region of m/z and mobility in a specified frame are fragmented with an elevated collision energy to produce MS2 spectra for that isolation region.

In complex samples, an intense precursor ion will likely have other precursor ions in its vicinity. If these nearby precursor ions are within the same band of mobility, they will be co-fragmented with the instrument-selected precursor ion and thus generate chimeric fragment spectra that can be used for additional peptide identification, albeit with more difficulty (12).

In our current implementation for data-dependant acquisition, the isolation windows are used to seed regions in which to look for peptide features, as features outside this region will remain unidentifiable. To form a window in the m/z and mobility plane for each precursor, the width in m/z and the begin and end of the scan dimension of its isolation windows is extended in each dimension – by the isolation window’s scan breadth in the mobility dimension to allow for determination of the precursor’s apex, and by 1 Da in the m/z dimension in case the monoisotopic peak for the precursor was not contained by the isolation windows, as shown in Figure 2. From this extended window in the m/z and mobility plane, a cuboid is formed by extending the window in the retention time dimension. The frames from the precursor’s isolation windows are extended forward and back in time by the user-specified base peak width (a characteristic of the chromatography gradient and configured from empirical observation) to ensure the precursor ion’s apex in retention time can be determined.

**Figure 2.**
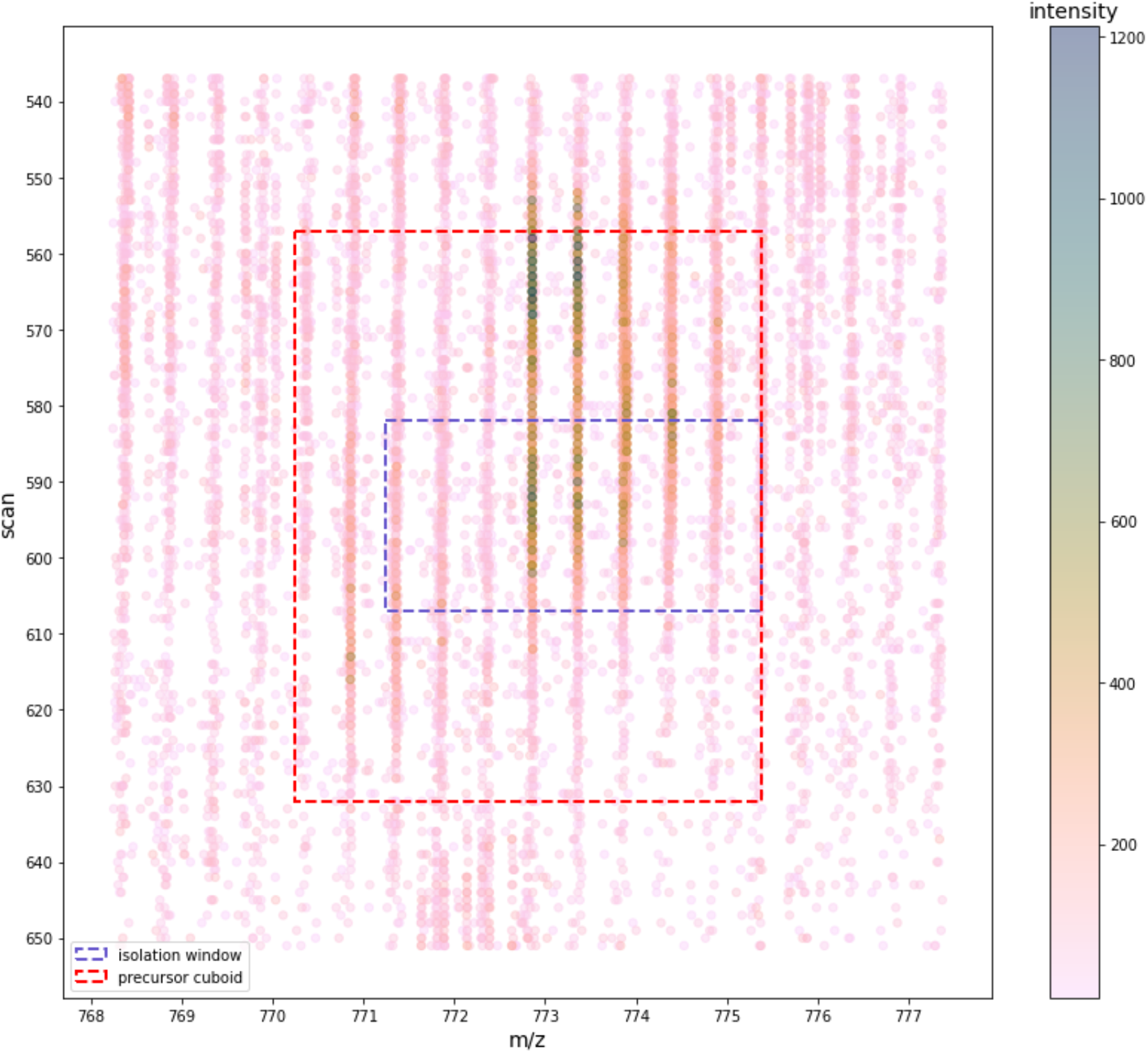
Formation of the precursor cuboid from the isolation windows recorded by the instrument for each precursor ion selected for fragmentation. Note the points shown are from all the frames in which this precursor was selected.

In the next section we describe how each precursor cuboid is resolved into one or more features, each with their constituent isotopic peaks.

### 4.2 Resolving spectra with intensity descent

The precursor cuboid defined with the approach described in 4.1 significantly reduces the peptide feature search space to a relatively tiny subset of the 4D raw data space. Within this reduced space however, more than one peptide feature may be present when two or more peptides have similar collisional cross section and m/z, and their elutions overlap in retention time. It is known that whether a peptide coelutes with other peptides has a greater effect on the likelihood of its identification than other attributes, such as its abundance (13). To determine the peptide’s identification, it’s crucial that the raw points within the cuboid be resolved to the isotopic peaks of the one or more precursors it might contain.

Figure 3A shows an example of the raw intensity readings from a precursor cuboid in the m/z dimension. For each ion there are many individual intensity readings; to facilitate feature deconvolution, these individual readings must be resolved to a single m/z value for each ion and its intensity.

**Figure 3.**
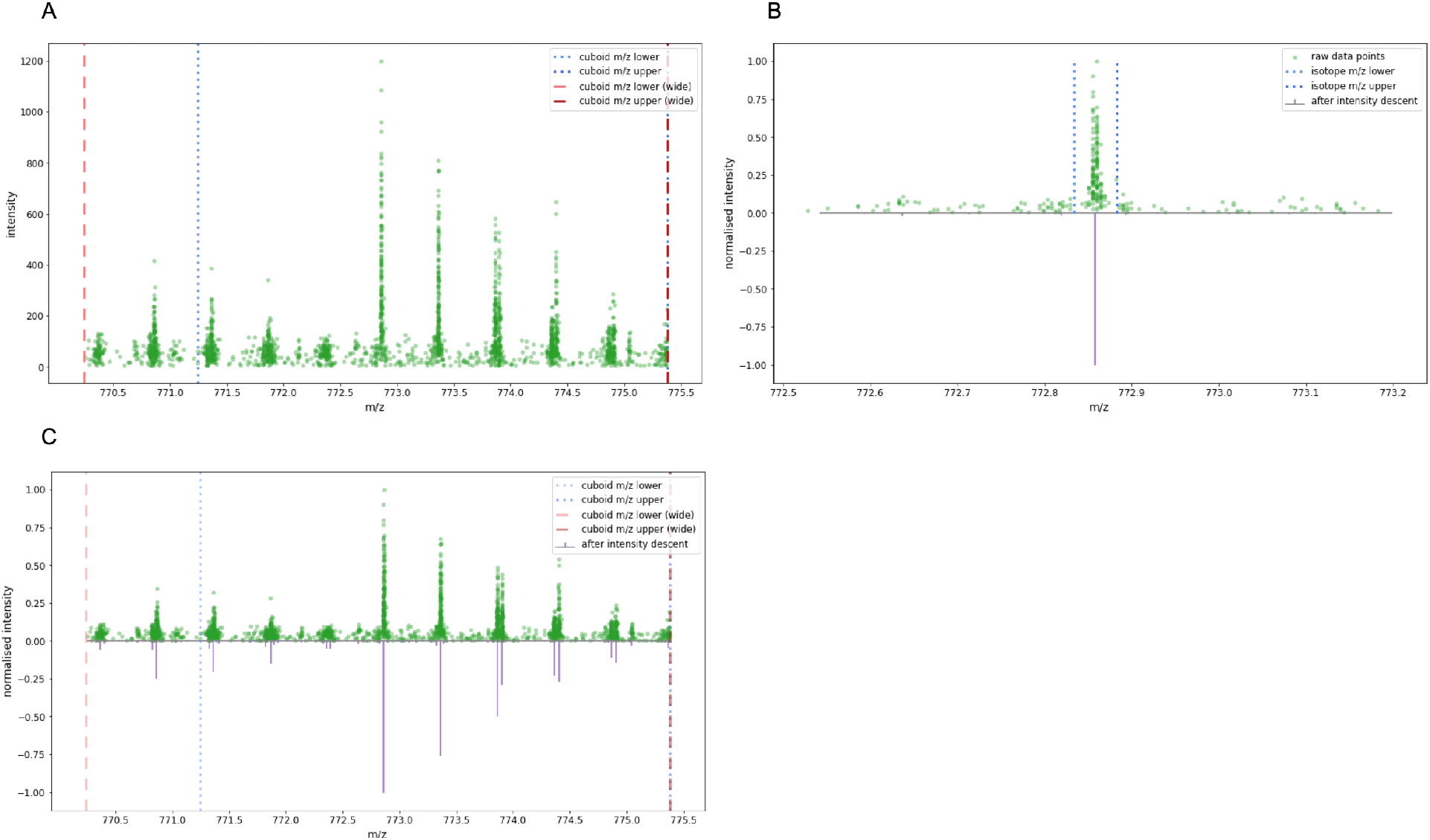
Resolving raw points to isotopic peaks. (A) Raw MS1 spectra within a precursor cuboid defined by a range of m/z, RT, and collisional cross-section values. (B) Comparing ms1 spectra before and after intensity descent for a selected peak within the precursor cuboid. The 3 σ bounds on each side are shown; the raw points within these bounds were summed and centroid with intensity weighting to form the single peak shown underneath. (C) Comparing the complete ms1 spectra for a selected precursor cuboid, before and after intensity descent.

In the m/z dimension, the intensity readings for each isotope of an ion form a Gaussian distribution (14). To deconvolve them, the spectra are scored by their fit to theoretical isotopic patterns for assumed charge states to find a probabilistic match. This task is simplified if the spectra is resolved to its significant ions, in a process called peak detection.

Approaches that use the first-and second-derivative zero crossings of smoothed raw data (15), and Kalman filters (16) to extract ion chromatograms have proven successful. More recently, the centWave (17) is a peak detection algorithm that uses a continuous wavelet transform (18,19) (CVT) applied to regions of interest. These approaches analyse ion intensity peaks in the m/z dimension through retention time, however they do not cater for the additional dimension of mobility.

To determine a single pair of m/z and intensity values for each precursor ion and its isotopes, a common approach is to create bins of fixed m/z width, find the intensity-weighted centroid of each bin, and sum the bin’s intensity. The main problem with this approach is it creates arbitrary m/z boundaries; the risk for intensity readings to be incorrectly attributed to an ion, or for an ion’s intensity to be split across a bin boundary is high (20).

As our objective is to perform feature segmentation in m/z, retention time, and mobility without eroding the instrument’s precision, we chose to develop a fast and simple peak detection algorithm based on intensity-seeded binning, which we call ‘intensity descent’. The approach uses the intensity of each ion as a guide to determine which intensity readings belong together in the same peak. The idea is similar to the ‘bucketing’ approach described in (15).

First, the algorithm looks for the most intense point within the spectral region of interest (in this case, the precursor cuboid’s extent in the m/z dimension) and gathers the points within a fixed window either side, finding the intensity-weighted centroid in the m/z dimension and summing the intensity values to determine the total intensity for the ion. The window width is determined at a particular m/z by calculating three standard deviations of a Gaussian distribution based on the instrument’s resolution (for the timsTOF we use 40,000 (5)), using the full width at half magnitude (FWHM) method.

~~~
# find 3sigma for a specified m/z
def calculate_peak_delta(mz):
   delta_m = mz / INSTRUMENT_RESOLUTION # FWHM of the peak
   sigma = delta_m / 2.35482 # std dev is FWHM / 2.35482
   peak_delta = 3 * sigma # 99.7% of values fall within +/-3 sigma
   return peak_delta
~~~

Having been summed and centroided, the points within the window are removed from the spectra, and the next most intense point is found. The process repeats until there are no points remaining to process in the region of interest. Figure 3B shows an example of a single peak within the precursor cuboid’s spectra and the raw points that were gathered to form it, and Figure 3C shows the spectra before and after intensity descent for the whole precursor cuboid.

To resolve the simplified peaks into series of isotopic peaks for candidate features, the precursor cuboid’s simplified spectrum is presented to the deconvolution algorithm in the Python `ms_deisotope` package (21). This function returns a set of proposed features derived by taking peaks as candidate isotopic peaks for a particular charge state, along with a score that indicates the quality of the proposed isotopic peak series against the BRAIN theoretical model of tryptic peptides (22,23).

Each panel in Figure 4 shows a feature proposed for the precursor cuboid spectra. In each panel, the coloured rectangles show the envelope of the feature, encompassing the isotopes and their intensities. The feature’s monoisotopic peak, and its charge state, are highlighted. The features up to a configurable maximum number that were scored above a specified threshold (also configurable) are added to a list of features obtained from all the precursor cuboids in the run to be rendered as an MGF file and searched against a database of tryptic peptides.

**Figure 4.**
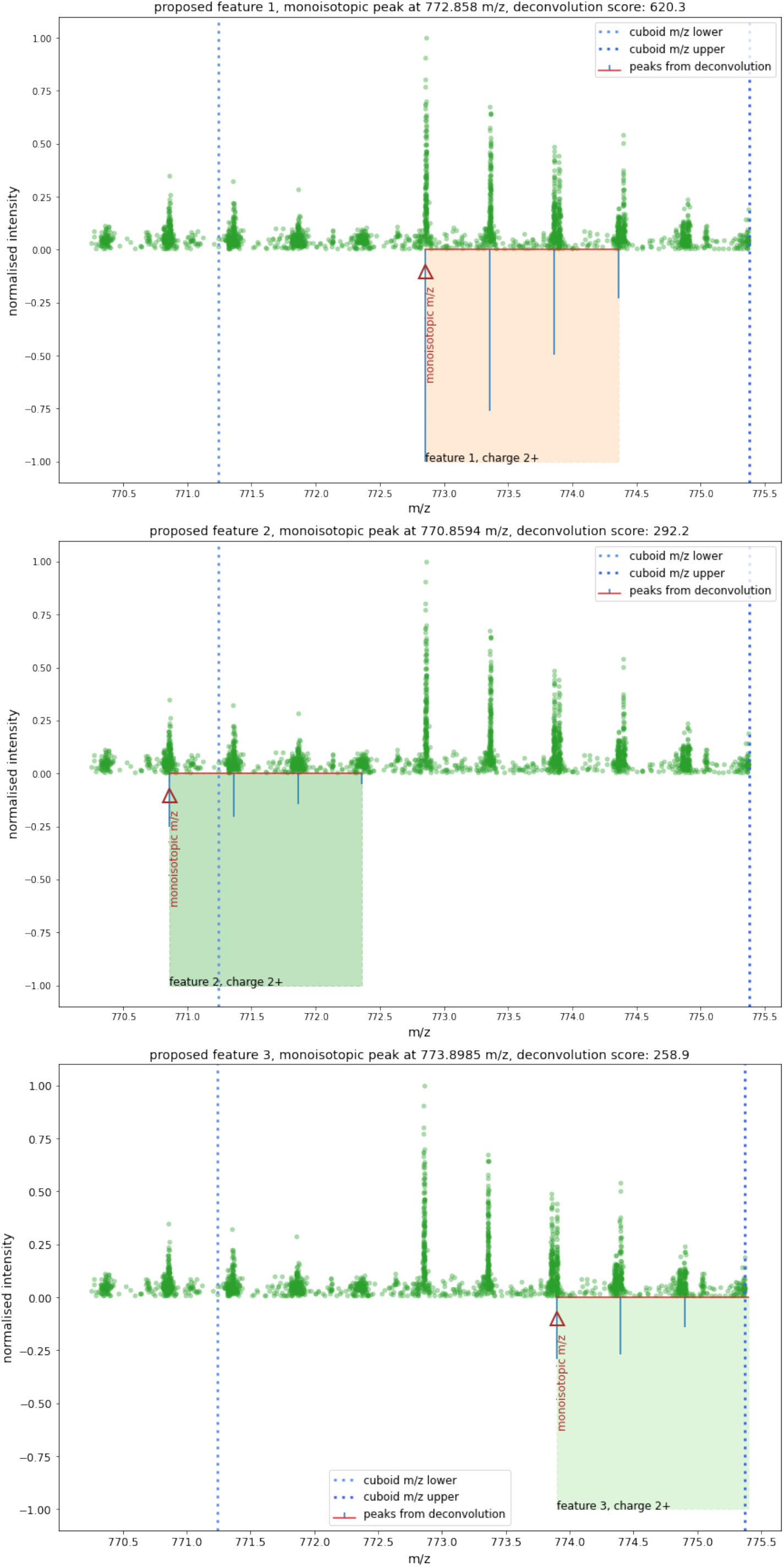
A set of proposed features derived from the simplified spectra for a precursor cuboid.

The monoisotopic peak of each proposed feature is analysed to find its extent in the mobility and retention time dimensions, by constraining the cuboid’s points in the m/z dimension to the peak’s ±3σ limits, as shown in Figure 5A.

**Figure 5.**
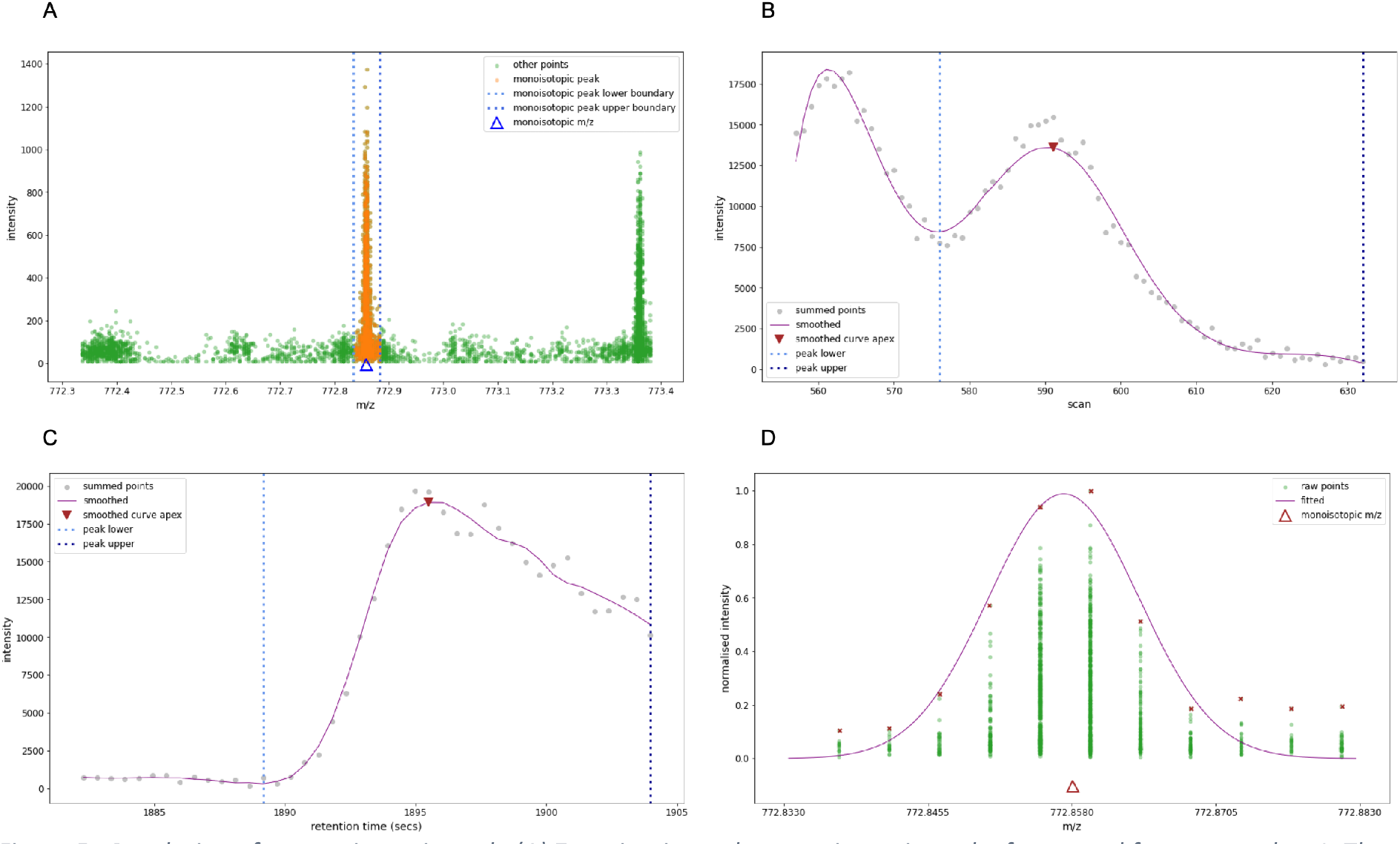
Resolution of a monoisotopic peak. (A) Zooming-in on the monoisotopic peak of proposed feature number 1. The dotted blue lines either side of the peak show the ±3 σ limits at the peak’s m/z. (B) The points in the monoisotopic peak collapsed to the mobility dimension. There are two peaks quite close together in the mobility dimension. The marked peak is the closest to the centre of the fragmentation event. The blue dotted lines show the valleys that have been determined on each side of the highlighted peak. (C) The points in the monoisotopic peak collapsed to the retention time dimension. (D) A zoomed-in view of the feature’s monoisotopic peak in the m/z dimension.

To determine the monoisotopic peak’s extent in the mobility dimension, we collect all its points and group them by their scan number and sum their intensity, flattening the points to the mobility dimension by summing the points that occur on the same scan. We chose a Savitzky-Golay filter (24) for its speed and effectiveness to smooth the points, and we use the peakutils Python package (25) to find the valley on either side of the peak, thus determining the monoisotopic peak’s extent in the mobility dimension, as shown in Figure 5B.

Having determined the extent of the monoisotopic peak in the m/z and mobility dimensions, the dimension-flattening procedure is repeated to determine the extent of the peak in retention time, as shown in Figure 5C.

The intensity of each isotope is calculated by summing the most intense point in the peak and the most intense point in the frame on either side. The intensity of the feature is the sum of the intensities of the first three isotopes. This method provides a robust measure of the feature’s intensity, as it is not determined by the intensity of a single isotopic peak.

We previously mentioned that the peak of an ion in the m/z dimension conforms to the curve of a Gaussian distribution. We confirm the monoisotopic peak’s Gaussian nature by fitting a Gaussian curve to the points of the peak in the m/z dimension and calculate the R-squared deviation of the observed and expected distribution. For this peak we can see in Figure 5D the peak is indeed a close fit to a Gaussian distribution, with an R-squared value of 0.9 compared to a Gaussian distribution.

On completion of these steps, the MS1 attributes of the feature required for searching against a peptide library have been determined: the monoisotopic m/z, the charge state, the feature’s intensity, and its apex and extent in both mobility and retention time. These attributes are included in the MGF’s header for each feature.

In this section we have introduced a method for resolving the isotopic peaks and determining some key attributes for the identification of a peptide feature contained in the precursor cuboids defined in 4.1. By constraining the challenge of feature detection in the vastness of the raw data space to thousands of much smaller precursor cuboids that can be processed in parallel, the complexity of segmenting individual peptides and their isotopic peaks is significantly reduced, thus increasing the confidence in the feature attributes presented to the subsequent peptide identification steps.

### 4.3 Correcting intensity readings in saturation

The timsTOF uses a high-gain electron multiplier called a microchannel plate (MCP) detector to count ions striking it on their descent from the top of the flight tube. MCP detectors produce a current of electrons when an incident ion forces zero or more electrons in the surface of the plate to be emitted and cascaded into subsequent plates in a multiplying effect (26). A 10-bit, 5×10^9^ samples/second analog-to-digital converter (ADC) (27) converts the current produced by the MCP detector to determine a measure of ion intensity, a proxy for relative ion abundance.

An MCP detector operating at high gain can appear to be saturated when many ions strike it in a short time interval. The electrons being ejected do not have sufficient time to be replenished before the next incident ion arrives. This can occur when the ion density is high, such as for ions in high abundance or ions in close proximity (28). The effect of saturation is that dynamic range for these high abundance ions is reduced.

Detector saturation is common in many types of mass spectrometer design, including TOF instruments (29). To address this situation in the timsTOF, a mass-dependent correction with base points and linear interpolation between the base points is used. The default corrections were determined by Bruker experimentally but are adjustable in the instrument configuration.

The effects of detector saturation can be observed by plotting the raw intensity values by m/z, as shown in Figure 6A for a typical run of a Yeast-HeLa-E.coli mixture. The limits of the detector and the instrument’s nonlinear discrete extrapolation of intensity readings are visible. In Figure 6C the empirical distribution function of the raw intensity shows the nonlinearities when intensity is higher than about 3000. Figure 6B shows an enlarged view of intensities around the 3000 mark. It is also evident that for the 550-1100 m/z range there is nonlinear intensity extrapolation between 2400 and 2600 intensity. The implication of detector saturation for peptide feature detection and the correct determination of isotopic peak height ratios is that some more intense points are in saturation and may not represent a true reading of the ion’s relative intensity. This in turn affects the calculation of the feature’s intensity and peptide quantification.

**Figure 6.**
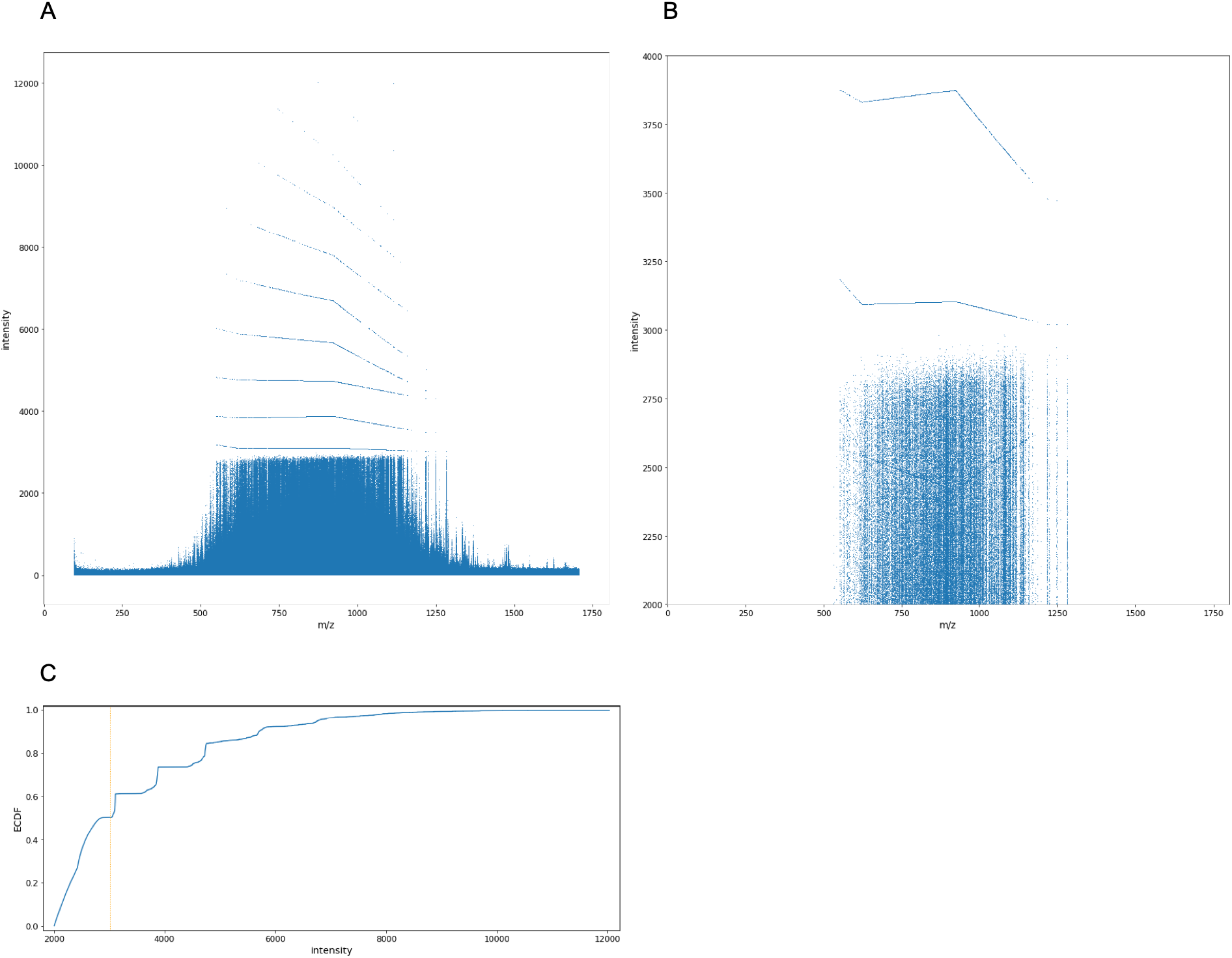
Effects of saturation on indicated intensity across the m/z range. (A) Raw data from analysis of the Yeast-HeLa-E.coli mixture, showing the nonlinearities in the readings where the detector is saturated. (B) Enlargement of the raw data intensity range. (C) The empirical cumulative distribution of the raw points, showing that more than half the raw points are affected by saturation.

Bilbao et al reported that detector saturation can affect mass accuracy and dynamic range, and they proposed an algorithm based on an averagine peptide model of isotopic peak intensities to adjust for the effect (30). Our approach to adjusting for detector saturation is to employ the Valkenborg theoretical model of peak height ratios for tryptic peptides (31) to improve the determination of the monoisotopic peak’s intensity when it is comprised of points that are in saturation. We have taken the raw intensity value of 3000 as an indication that a reading was in saturation. When isotope 0 (i.e. the monoisotopic peak) is comprised of at least one point above the saturation threshold, the model is used to infer what the intensity should be for a peptide of its monoisotopic mass, based on the intensity of the closest isotope in the series that does not include points in saturation. An example of this approach is shown in Figure 7, where the first four isotopic peaks comprised points in saturation. In this example, the intensity of the first isotopic peak that was not comprised of points in saturation (peak index 4) was used to infer the intensity of the next isotopic peak (peak index 3), and that peak’s intensity to infer the next peak, and so on until the monoisotopic peak is reached. The figure shows the difference between raw intensity including saturated readings, and the inferred intensity using the intensity of the first non-saturated isotopic peak.

**Figure 7.**
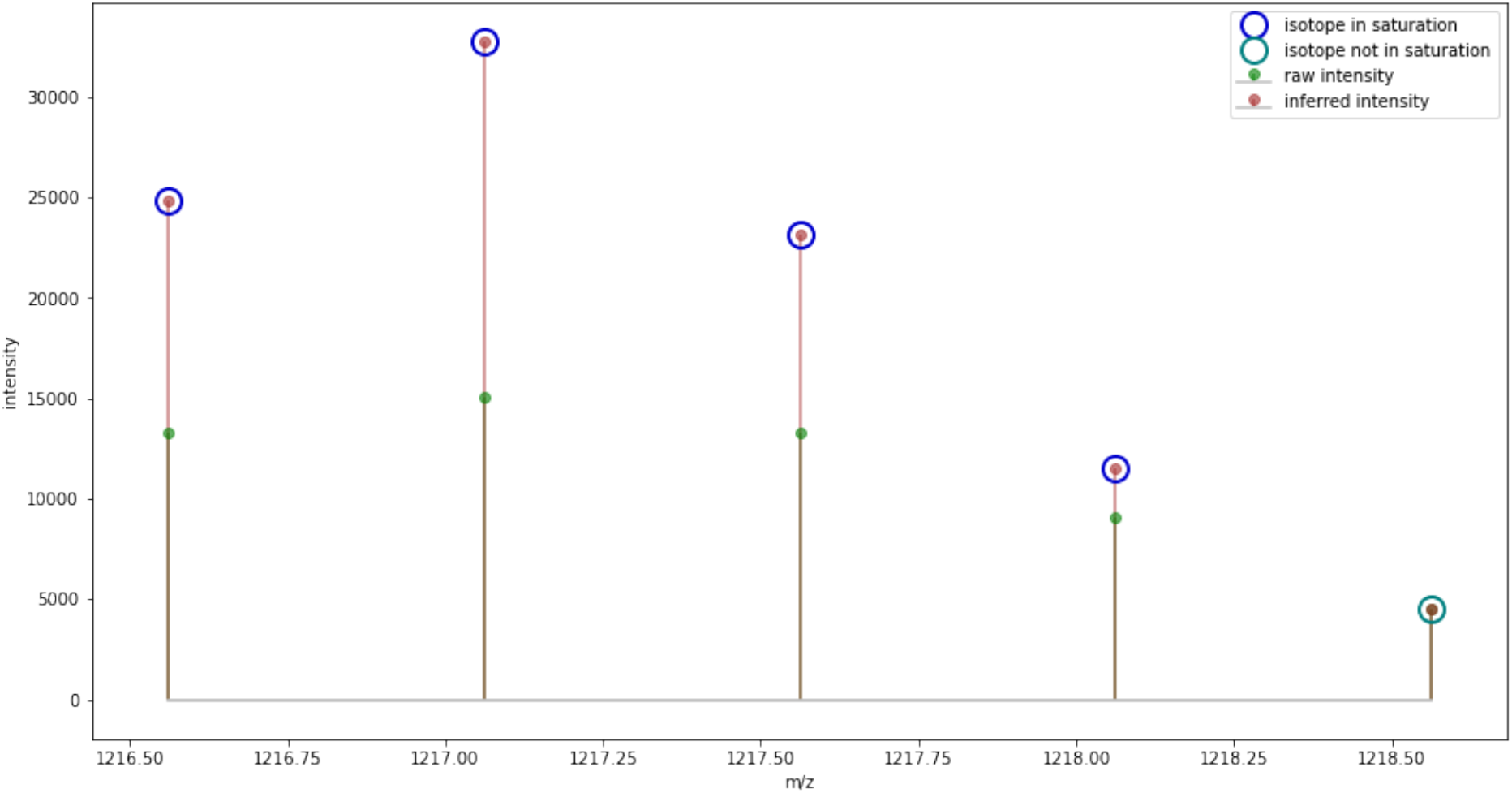
using the theoretical model of tryptic peptide peak height ratios to infer isotope intensities from the first non-saturated isotope

We have found that monoisotopic peak intensity saturation affects about 5% of identified peptides in 200ng samples of Yeast-HeLa-E.coli mixture. Of all the detected features where the monoisotopic peak was in saturation, the third isotope (index 2) was most commonly used to infer the intensity (the distribution is shown in Figure 8). This might be explained by the intensity of second isotopic peak (index 1) usually being of similar or greater intensity than the monoisotopic peak. Therefore, if the monoisotopic peak is in saturation, the second isotopic peak will usually also be in saturation.

**Figure 8.**
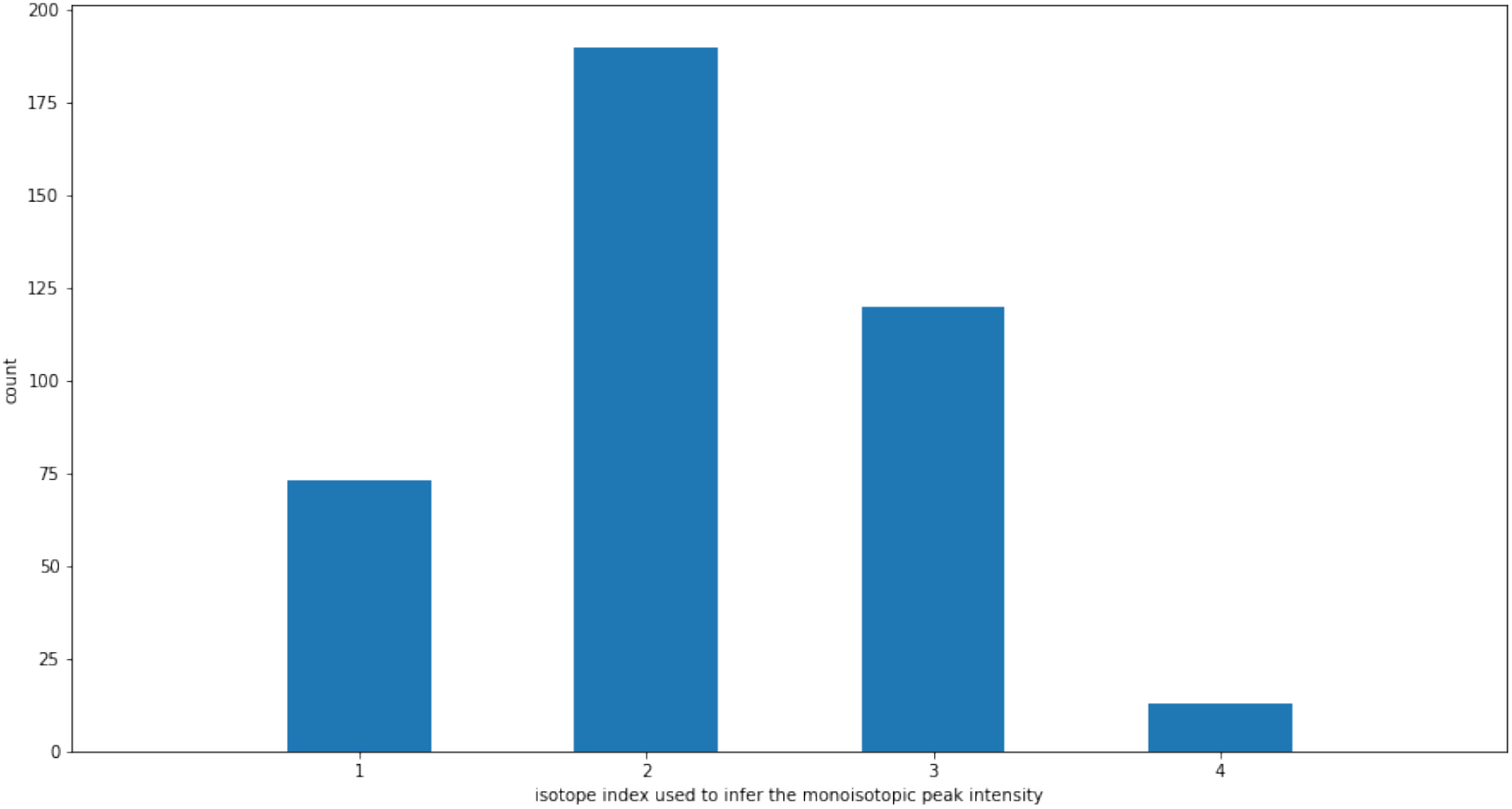
the distribution of isotope index used to infer the intensity of the monoisotopic peak

When isotopic peak intensity is calculated by taking the top three points nearest the apex of the feature’s peak in retention time, the maximum intensity of a peak without saturation is 3 × 3000 = 9000. The ranges of isotopic peak height ratios predicted by the theoretical model for tryptic peptides vary according to the isotope index and the peptide’s monoisotopic mass; the model does not predict peak height ratios for all isotopes across the same range of monoisotopic mass. For example, as shown in Figure 9, the minimum monoisotopic mass range predicted by the model for the seventh isotope (index = 6) is 1550 Da. Therefore, when the seventh isotope is the first unsaturated isotope, the model can only be applied to peptides with monoisotopic mass greater than 1550 Da.

**Figure 9.**
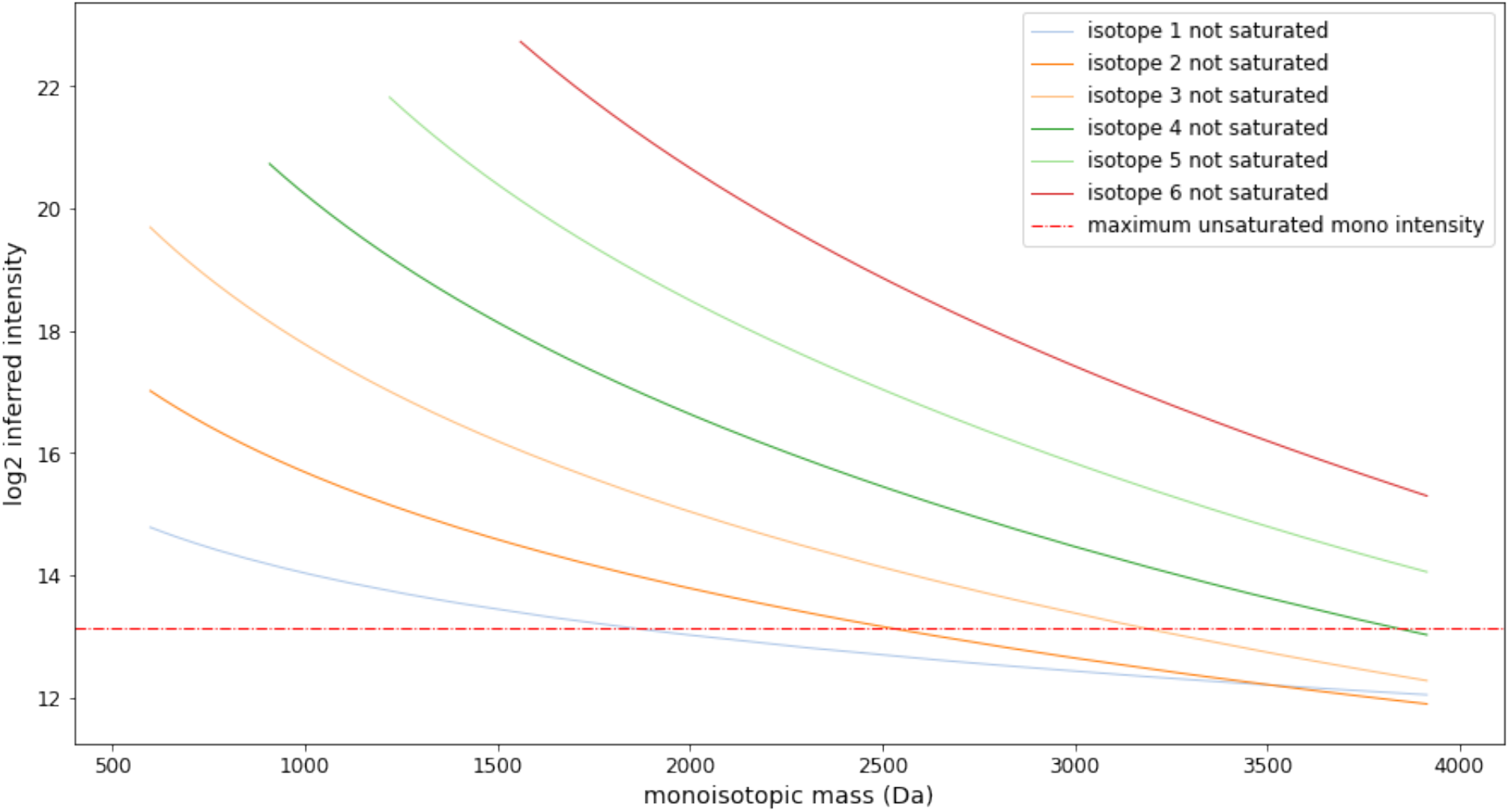
the maximum inferred intensity of the monoisotopic peak depending on which isotope is used to infer its intensity

In the Yeast/HeLa/E.coli mixtures analysed, only 0.03% of features had a monoisotopic peak that could not be inferred because all the detected isotopes had points in saturation, and 0.01% of features could not be inferred because of their monoisotopic mass and the number of their isotopes that included points in saturation were outside the model’s applicable range. For higher load samples, where the detector is overloaded to a much greater extent, many more features would have a monoisotopic peak that contains points in saturation. Our approach to correcting the monoisotopic peak’s intensity by inference could facilitate the recovery of highly saturated data.

Consistent with findings reported by Bilbao (30), in the example shown, we observed a modest improvement of dynamic range in the identified features that had a monoisotopic peak with at least one point in saturation and then corrected, as shown in Figure 10. Correcting for saturation improved the dynamic range of features, increasing from 0.97 orders of magnitude without correction to 1.14 orders of magnitude with correction.

**Figure 10.**
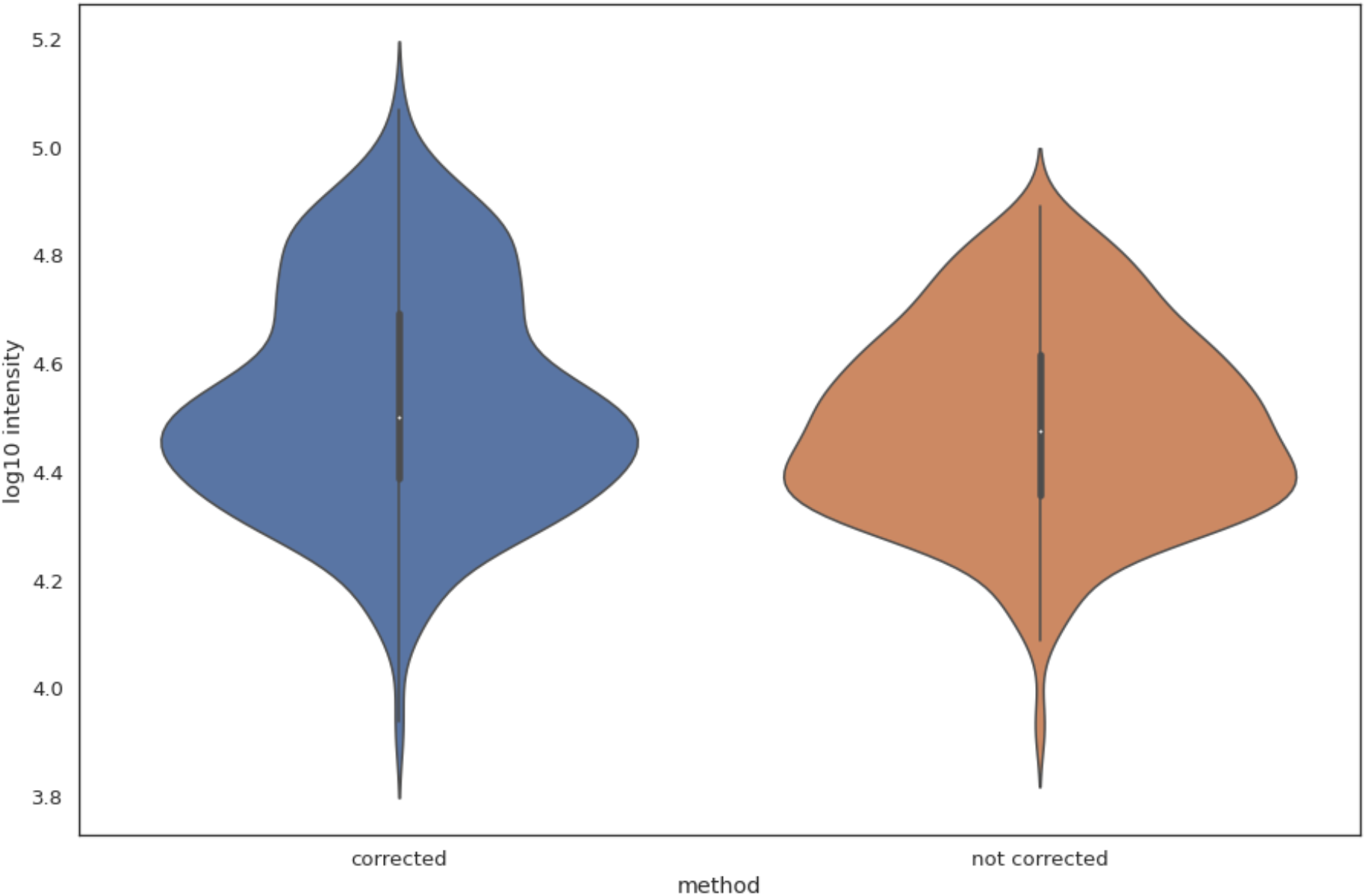
dynamic range of identified features with their monoisotopic peak in saturation with and without correction for saturation

In addition to points in saturation affecting the calculation of a monoisotopic peak’s intensity, Bilbao (30) reported that correction for intensity saturation also improved mass accuracy. To gauge the effect of intensity saturation on mass accuracy in timsTOF data, we removed points in saturation prior to intensity-weighted centroiding the m/z of isotopes, and found that mass error of identified peptides slightly increased (Figure 11) by an average of 0.064 ppm.

**Figure 11.**
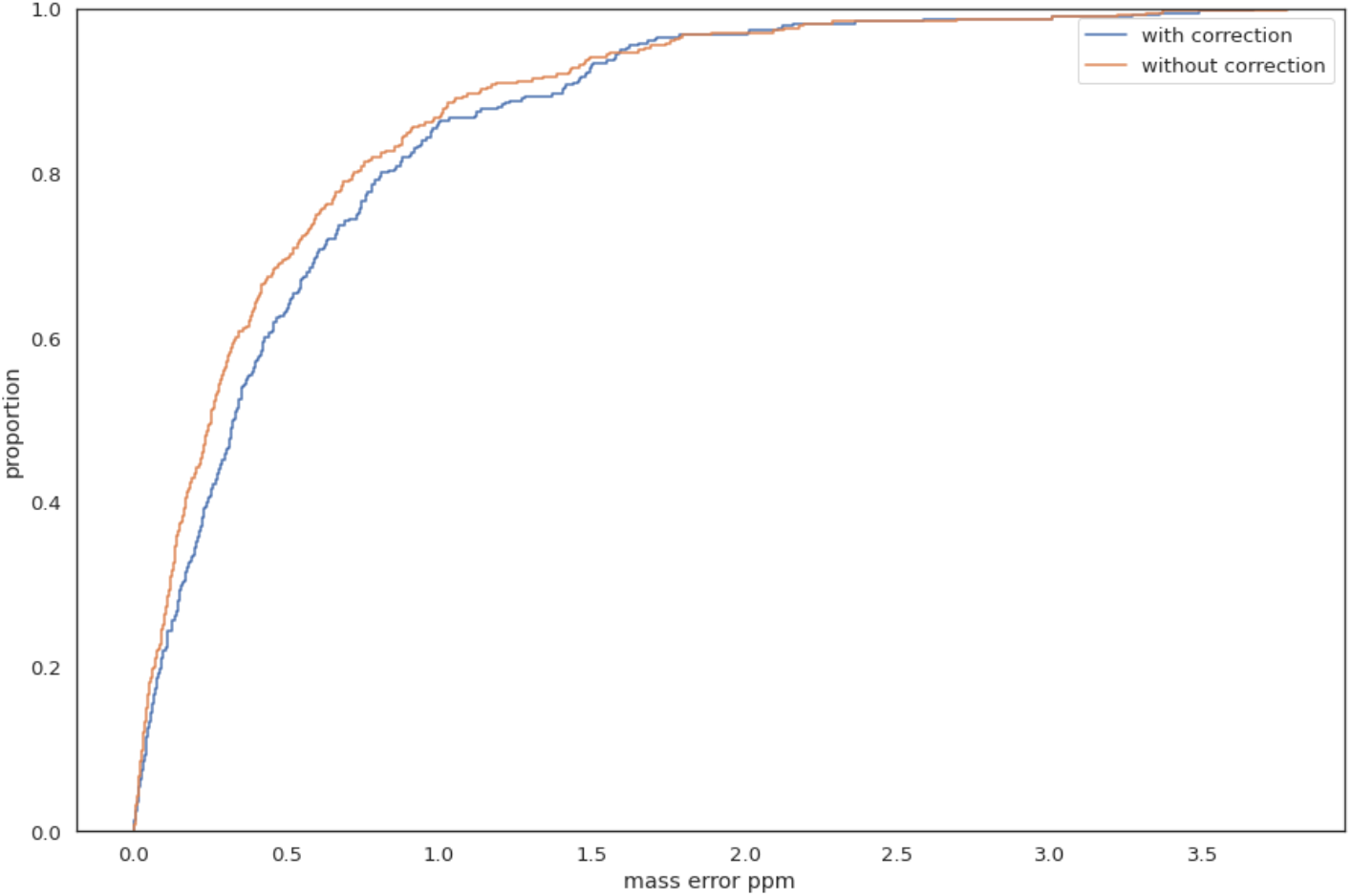
ECDF of mass error for identified peptides with and without correction for saturation

We conclude that the instrument’s extrapolation of intensity values for peaks in saturation is effectively correcting shifts in m/z due to saturation.

To determine the effect correcting for detector saturation on peptide quantification, we analysed ten technical replicates of a Yeast and UPS2 mixture, and compared the mean detected protein concentration with ten technical replicates of Yeast and UPS1 mixture. The UPS1 mixture comprises 48 human proteins of diverse molecular weight and present in the same concentration of 5 pmoles of each protein. The UPS2 mixture (32) comprises the same 48 human proteins, arranged in six groups of different concentrations, ranging from 50,000 fmol to 0.5 fmol of each protein. When we compare the ratios of observed protein quantity in UPS2 relative to its quantity in UPS1, we should expect to see an improvement in the correlation between the observed protein ratios and the expected protein ratios when intensity correction is applied to peptides in saturation. In the UPS2 sample we observed 17.7% of monoisotopic peaks were affected by saturation, while in the UPS1 sample 9.2% of monoisotopic peaks were affected. The difference of affected monoisotopic peaks is expected because of the varying concentrations in the UPS2 experiment. The Top3 method as described in (33) was used to determine protein abundance. It was observed that 64.1% of the proteins identified in the UPS2 experiment had at least one of their top-3 peptides adjusted for saturation, while for the UPS1 experiment 47.7% of the proteins identified had their top-3 peptides adjusted. This means that correction for saturation plays a significant role in protein quantification determination with the Top3 method, as the most intense peptides used for the calculation are most likely to have monoisotopic peaks comprising saturated points.

The same experiments were analysed with MaxQuant (Version 1.6.17.0). To compare our feature detection with MaxQuant’s, the features detected by MaxQuant as stored in its APL (Andromeda peak lists) files were rendered as MGF files and searched with Comet and Percolator against the same Yeast-UPS1-UPS2 FASTA database that we used to search our MGFs. The ratios of abundance for the proteins found in common by our method and MaxQuant in both UPS1 and UPS2 samples are shown in Figure 12. For proteins with the same concentration in UPS1 and UPS2 and therefore having an expected log10 ratio of zero, no bar is shown. For those proteins that are not affected by saturation, the bars for ‘with saturation correction’ and ‘without saturation correction’ are the same height.

**Figure 12.**
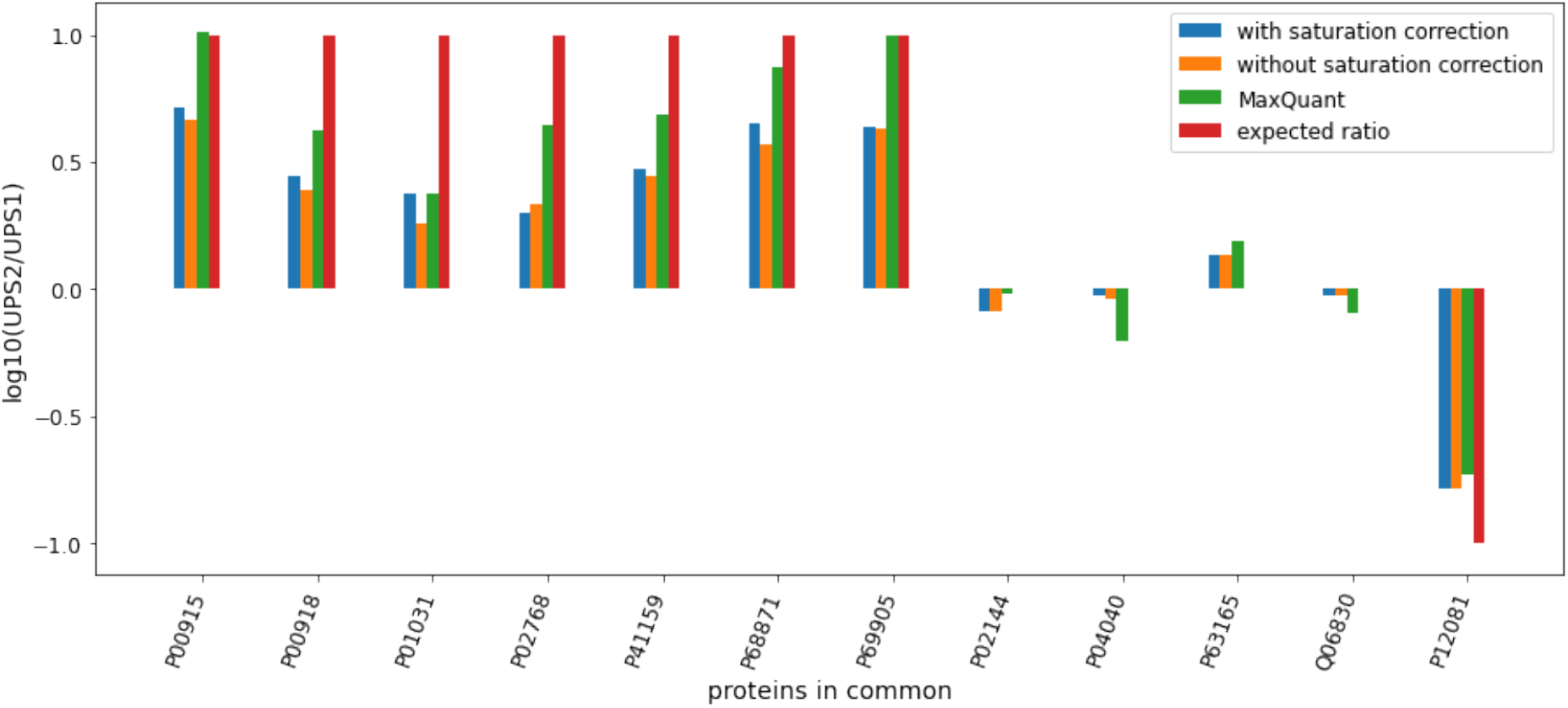
comparison of intensity ratios for proteins detected in the Yeast+UPS1 and Yeast+UPS2 experiments by our method and MaxQuant. A protein abundance ratio is shown if it meets all criteria for comparison: (i) at least three peptides for the protein were found, (ii) the protein was found by our method in at least one technical replicate of UPS1 and at least one of UPS2, (iii) the protein was found by MaxQuant in at least one technical replicate of UPS1 and at least one of UPS2, and (iv) the protein was found by both our method and MaxQuant.

For proteins with monoisotopic peaks affected by saturation, in most cases our approach to correct the intensity yields a better alignment with the expected abundance ratio between UPS2 and UPS1. For the proteins of 50 pmoles concentration in UPS2, MaxQuant had a better alignment with the expected ratio. For the proteins with 5 pmoles in UPS2, our approach had better alignment with the expected ratio than MaxQuant for three of the four proteins (P04040, P63165, QO6830). For the protein P12081 with 0.5 pmoles in UPS2, our ratio was more accurate than MaxQuant.

In the previous two sections we have focussed on methods we have developed to determine the attributes of precursor ions in MS1 spectra. Another critical step in the identification of peptides is the resolution and characterisation of their fragment ions, as it is by analysing the components of a peptide produced by fragmentation that we gain confidence in the identification. In the next section, we discuss a method we developed to simplify complex fragment MS2 spectra.

### 4.4 Simplifying ms2 spectra with mass defect windows

Collision-induced fragment ions result from the cleavage at amide bonds when the precursor ions collide with molecules of the non-reactive gas. The fragment ion spectra in the timsTOF are generally more complex than lower-energy collision-induced disassociation (CID) mass spectrometers (34), and this fact provides motivation for exploring ways to simplify the spectra using prior knowledge.

It is known that all isotopes have a mass defect, the phenomenon where each isotope releases different binding energy when it forms its nucleus (35). The mass defect of a molecule is defined as the difference between its exact mass (i.e., the sum of the atomic masses of constituent atoms) and its integer mass (i.e., the sum of the integer masses of the constituent atoms) (36). Relative to the reference mass of Carbon-12, the mass defect may be positive or negative.

Mass defects have many useful applications, particularly in the analysis of high resolution mass spectra (37). Mann *et. al*. first quantified the phenomenon for mass spectrometry applications by calculating peptide mass from sequence databases for peptides up to 2 kDa (38), identifying areas where peptide masses must lie and areas where they cannot, which he labelled ‘forbidden zones’. More recent analysis reported that mass defects observed in tryptic peptides and their modifications are slightly narrower than earlier described (39,40).

Nefedov et al (41) extended the idea and computed reference tables for all theoretically possible tryptic peptides up to 3 kDa at a resolution of 0.001 Da. They found gaps in the mass dimension in which peptides cannot fall, and ‘quiet’ regions where a peptide rarely falls. This knowledge can be used in a simple way to reduce the complexity of MS2 spectra and help to facilitate fragment ion extraction through the reduction of the number of points. By knowing that these points are likely chemical or electronic noise and not signal from peptides, we can remove those that exist in the gaps between mass defect windows. We used this phenomenon to simplify the fragment spectra for each feature in the MGF we generated for the features proposed in the deconvolution step. Although mass defect windows can also be applied to MS1 data, the most benefit is gained when using them to filter MS2 data; the task of fragment ion deconvolution is much more challenging due to the presence of co-fragmented peptides and noise.

To resolve fragment spectra, the MS2 raw points contained by the bounds of the fragmentation event in the mobility and retention time dimensions were extracted (Figure 13). In the m/z dimension we did not apply bounds, as the m/z of a fragment ion may be below or above its precursor ion. Using the same procedure as for MS1 ion resolution, we collapsed these raw MS2 points to the m/z dimension and performed intensity descent to simplify the spectra.

**Figure 13.**
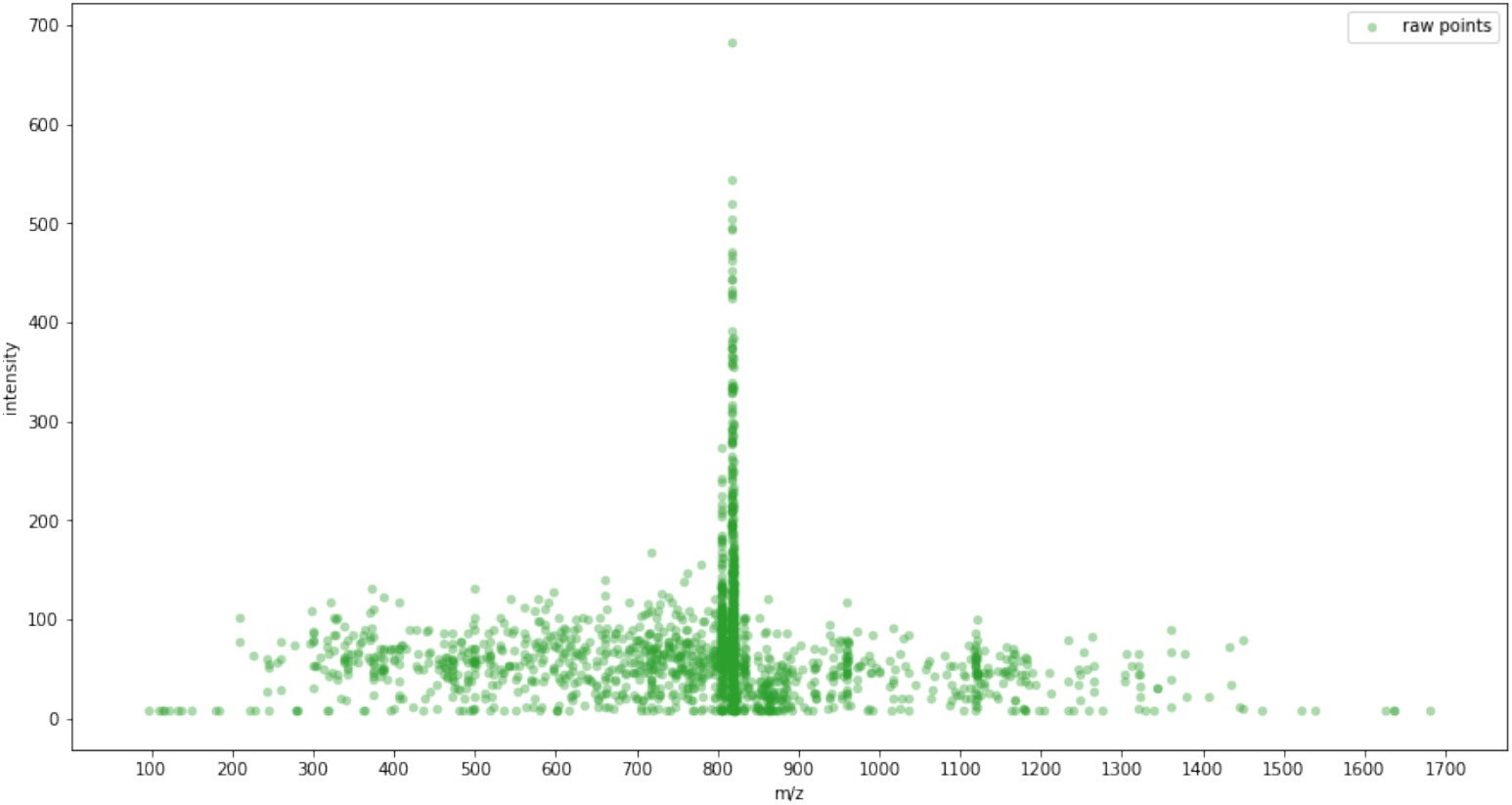
the raw MS2 points associated with an example precursor cuboid

As for MS1 spectra, we used the `deconvolute_peaks` function from the `ms_deisotope` package but with different settings appropriate to MS2 deconvolution to propose likely features and determine their monoisotopic m/z, isotopes and charge state from the theoretical model of tryptic peptides. The fragment ions in the MGF expected by crux’s comet function is the singly protonated mass, so to the neutral mass from the deconvolution step the Hydrogen proton mass of 1.00727647 Da was added. To apply filtering by mass defect windows, a fragment ion was removed if its neutral mass did not sit within the bounds of a mass defect window. In this sample a 21% reduction in fragment ions was observed when filtering was applied.

The mass defect windows were implemented in Python as bins of mass ranges, generated as follows:

~~~
def generate_mass_defect_windows(mass_defect_window_da_min, mass_defect_window_da_max):
 bin_edges_l = []
 for nominal_mass in range(mass_defect_window_da_min, mass_defect_window_da_max):
    mass_centre = nominal_mass * 1.00048
    width = 0.19 + (0.0001 * nominal_mass)
    lower_mass = mass_centre - (width / 2)
    upper_mass = mass_centre + (width / 2)
    bin_edges_l.append((lower_mass, upper_mass))
 return bin_edges_l
~~~

Each fragment ion was assigned to a bin according to its neutral mass. Those ions that were not assigned to a bin were removed. Even intense ions can be removed in this process, as can be seen in Figure 14, which shows a magnified portion of the neutral mass axis for the fragment ions before and after filtering by mass defect windows.

**Figure 14.**
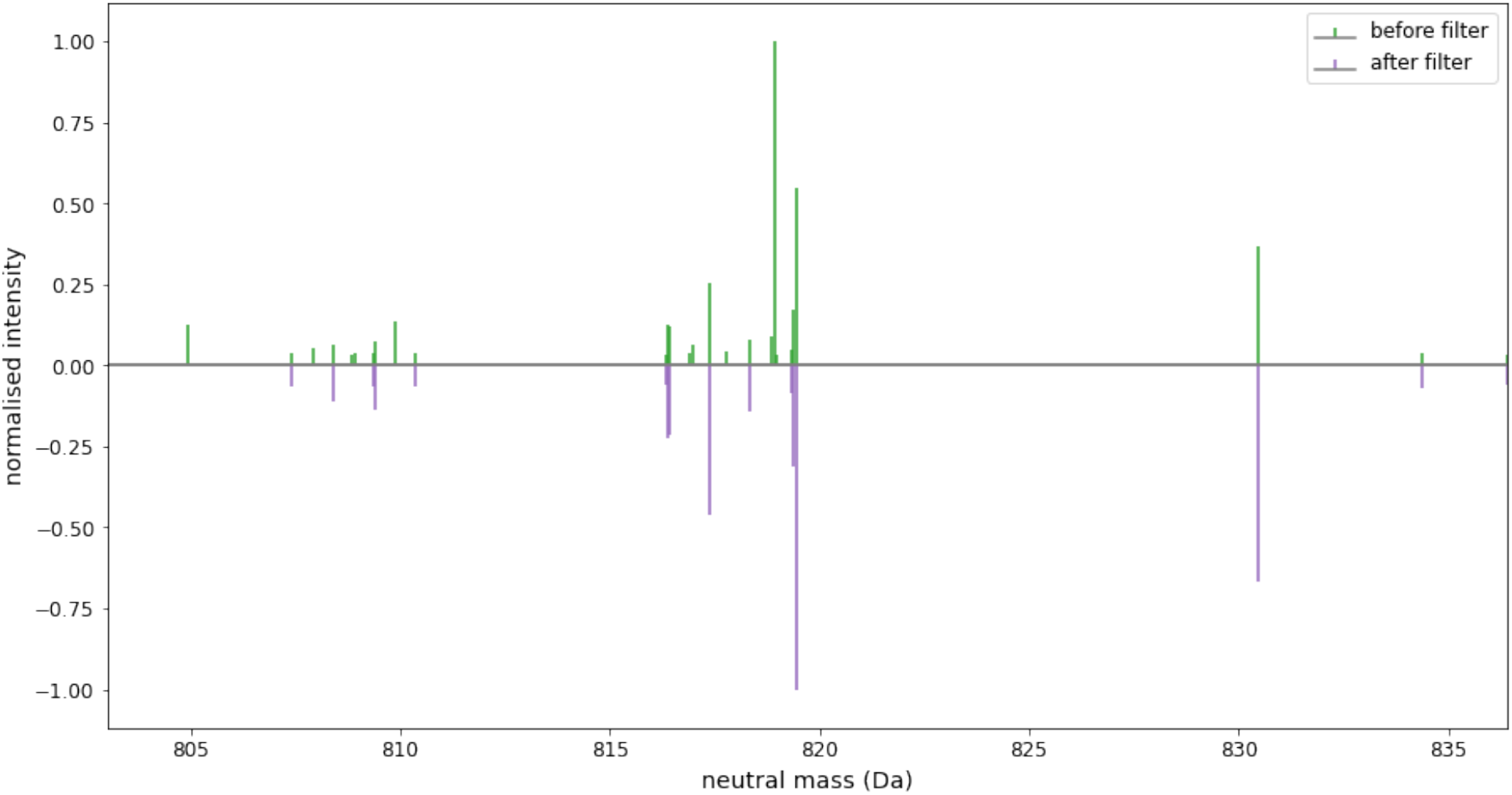
an enlarged segment of the neutral mass axis showing some examples of fragment ions being removed if they did not occupy a mass defect window

To assess whether more confident identifications were achieved with mass defect window filtering applied to the fragment ions of each feature, two MGF files were prepared for the same features detected in a run. One MGF has mass defect window filtering applied to the fragment ions, and the other MGF did not. Both MGFs were searched against a FASTA database with crux comet and percolator, and the score from percolator was compared for each identification in common. Identifications with q-value greater than 0.01 were removed to reflect a 1% FDR. A significant time reduction for rendering or search of the MGF was not observed when mass defect window filtering was applied, nor was there an increase in the number of unique peptides identified. However, of the peptide sequences identified in both MGF files, Figure 15 shows the identification is more confident when the fragment ion list is simplified by removing ions that are not within a mass defect window. For peptide sequences identified both with and without mass defect window filtering, an average 10.4% improvement in percolator score was observed with filtering applied. As the filtering is implemented in a few lines of Python code with little computational cost, we believe the benefit in peptide identifications clearly makes it a worthwhile inclusion in our processing steps.

**Figure 15.**
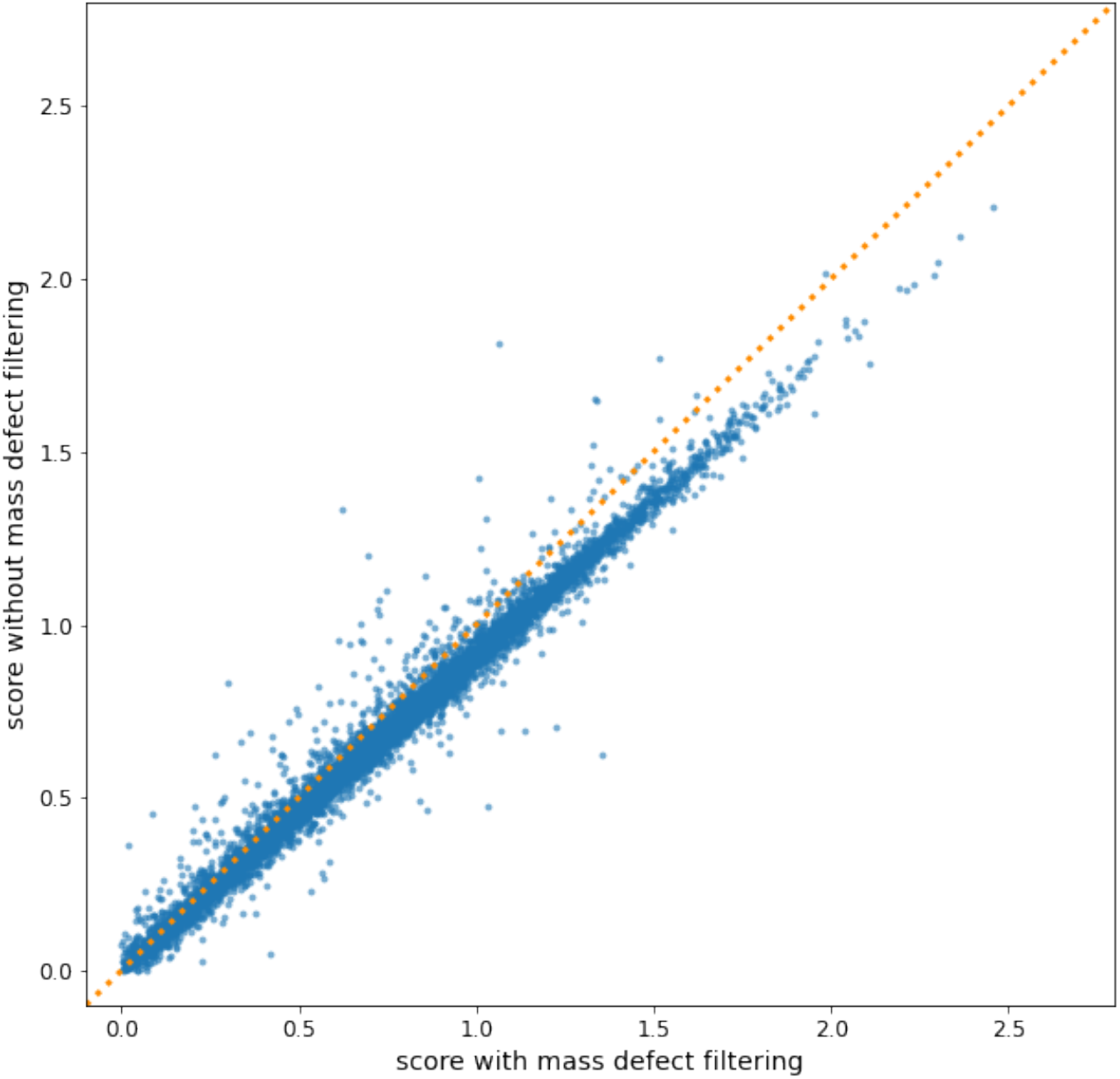
comparison of the score from percolator for identified features with and without mass defect window filtering

In the previous sections we have presented techniques for resolving and simplifying MS1 and MS2 spectra to improve the quality of peptide features. In the next section we discuss how applying these techniques in combination can achieve higher quality peptide identifications.

### 4.5 Comparing feature detection with MaxQuant

To compare our feature detection approach with MaxQuant, we used MaxQuant version 1.6.17 to process the same sample. We converted MaxQuant’s APL files to an MGF file and used Crux Comet and Percolator with the same parameters and FASTA file that we used for our own pipeline. MaxQuant achieves a higher Percolator score on average than our identifications, suggesting that MaxQuant does a better job at resolving and deconvolving fragment ions (Figure 16A). Further work on our processing would involve tuning our fragment spectra deconvolution. However, for the features found in common, our mass accuracy is higher than identifications from MaxQuant (Figure 16B).

**Figure 16.**
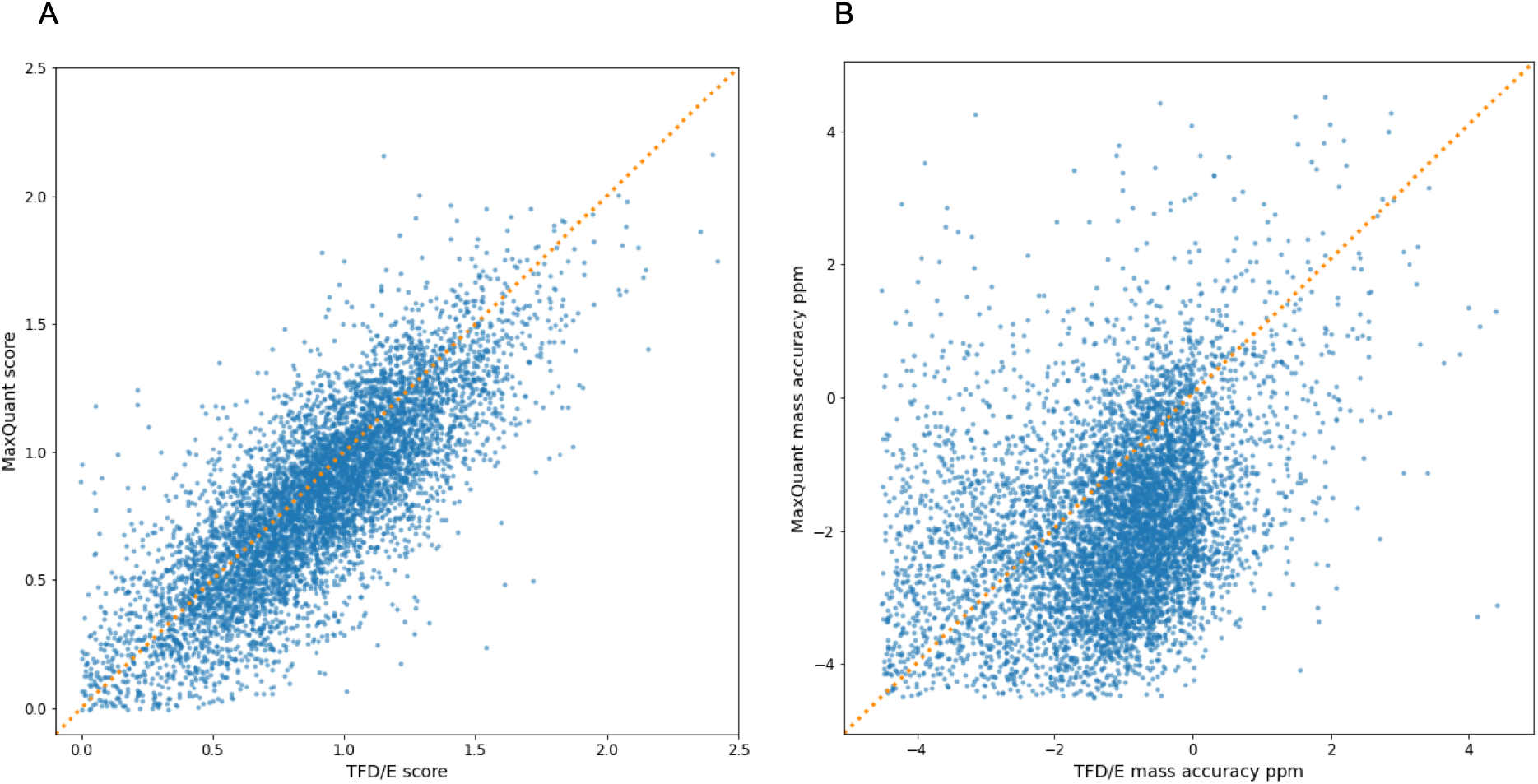
Comparing feature score and mass accuracy for features found by ours and MaxQuant. (A) The percolator score for peptide sequences identified in common. (B) The mass error ppm for the peptide sequences identified in common.

## 5 Discussion

In this work, we described our approach to the fundamental challenges in processing 4D timsTOF data: how do we find features, distinguish them from noise, deal with their signal complexity, and determine their monoisotopic peak. We have shown that intensity descent is an effective method to simplify spectra in MS1 and MS2, and how it considers the resolving power of the timsTOF. We have seen that when the instrument selects a precursor ion for fragmentation, there are often other viable features in the vicinity, often overlapping, and through deconvolution we can extract additional features apart from the instrument-selected precursor that are worthy of including in the database search of tryptic peptides.

Our approach for addressing the issue of detector saturation improves the dynamic range of the features extracted and yields a better alignment with the expected ratio of protein abundance in UPS1 and UPS2. Overall, this correction for intensity saturation of the monoisotopic peak in peptides translates to better quantification of proteins compared with not correcting.

Unlike the results reported by Bilbao on a Agilent 6224 TOF MS (30), we did not observe an improvement in the mass accuracy of peptide identifications by correcting for saturation, suggesting that the extrapolation performed by the timsTOF firmware when saturation occurs is dealing with the saturation effect on m/z readings without detriment to the mass accuracy.

We have built upon the work by Mann (38) and Nefedov (41) to show that MS2 spectra can be simplified through filtering of fragment ions using mass defect windows. Although using our test data we only found a small benefit of peptide identification confidence reported by Percolator, we believe that the small computational cost of the filtering makes it worthy of inclusion in the data processing.

Finally, we have shown that these approaches collectively result in an improvement in mass accuracy compared to MaxQuant. We believe this finding proves our original hypothesis: that these techniques in combination achieve a better peptide identification result than what is achieved without them.

A guiding principle in conducting this research was to embrace the ion mobility dimension provided by the timsTOF on an equal footing with the m/z and retention time dimensions, a feature we felt was neglected in other open-source processing tools for the timsTOF. Our algorithms achieve lower mass error, more dynamic range, and higher peptide identification confidence. By making available the source code in the popular and accessible Python language, and leveraging other open-source tools such as Comet and Percolator, our contribution is an experimental sandbox for feature detection. Our aim is to encourage experimentation with and improvement of these techniques, and to gain more insight into the detailed programmatic steps involved with identifying peptide sequences from raw timsTOF data.

## 6 Materials and Methods

### 6.1 Sample preparation

#### 6.1.1 Proteome Benchmark Dataset

Commercial tryptic digests of *S*.*cerevisiae* (Yeast, Promega, #V746A), human K562 cells (Promega, #V695A) and *E*.*coli* (MassPREP standard, Waters, #186003196) were reconstituted in 2% ACN/0.1% FA to final concentration of 0.1 µg/ul. To generate the hybrid proteome samples, purified peptides from each of the three species were combined in different proportions as previously described (42) and as follows: sample YHE211 consisted of 30% w/w Yeast (3 µg), 65% w/w Human (6.5 µg) and 5% w/w *E*.*coli* (0.5 µg); sample YHE114 consisted of 15% w/w Yeast (1.5 µg), 65% w/w Human (6.5 µg) and 20% w/w *E*.*coli* (2 µg); sample YHE010 consisted of 0% w/w Yeast (0 µg), 100% w/w Human (6.5 µg), and 0% w/w *E*.*coli* (0 µg). Ten replicates of each proteome mixture were subjected to LC-MS/MS analysis on a timsTOF Pro mass spectrometer.

#### 6.1.2 Dynamic Range Benchmark Dataset

UPS1 and UPS2 standards (Sigma-Aldrich) were combined with commercial intact Yeast protein (Promega, #V7341) by mixing 50 µg of Yeast protein with 3.2 µg of UPS1 or USP2 subjected to enzymatic digestion with Trypsin Gold (Promega, 1 µg) for overnight at 37 degrees Celsius using the FASP digestion method (43). Lyophilised peptides were reconstituted in 2% ACN and 0.1% FA. Ten replicates of each peptide mixture were subjected to LC-MS/MS analysis on a timsTOF Pro mass spectrometer.

#### 6.2 LC-MS methods

The digested proteome mixtures were separated by reverse-phase chromatography on a C18 fused silica column (i.d. 75 μm, o.d. 360 μm × 25 cm length, 1.6 μm C18 beads) packed into an emitter tip (IonOpticks, Australia) using a nanoflow HPLC (M-class, Waters). The HPLC was coupled to a timsTOF Pro mass spectrometer (Bruker Daltonics, Bremen) using a CaptiveSpray source. Peptides were loaded directly onto the column at a constant flow rate of 400 nL/min with buffer A (99.9% Milli-Q water, 0.1% FA) and eluted with a 20-minute linear gradient from 2% to 34% buffer B (99.9% ACN, 0.1% FA).

The timsTOF Pro was operated in PASEF mode using Compass Hystar 5.1 and otofControl settings were as follows: Mass Range 100 to 1700m/z, 1/K0 Start 0.85 V·s/cm^2^ End 1.3 V·s/cm^2^, Ramp time 100 ms, Lock Duty Cycle to 100%, Capillary Voltage 1600V, Dry Gas 3 l/min, Dry Temp 180°C, PASEF settings: 4 MS/MS scans (total cycle time 1.27sec), charge range 0-5, active exclusion for 0.4 min, Scheduling Target intensity 24000, Intensity threshold 2500.

### 6.3 Mass Spectrometry Analysis

The raw data was extracted from the instrument database and processed with our bespoke software for feature detection. For peptide identification and protein inference we used Crux Comet and Percolator. The mass tolerance for the initial search was 20 ppm; the search following mass recalibration was 4.5 ppm. Settings were a maximum of 2 missed cleavages, a bin tolerance of 0.02 Da for fragment ions, a bin offset of 0, and a default peak shape. The MH+ peptide mass range for analysis was 700-5000 Da.

The FASTA database was created for the Yeast/HeLa/E.coli proteome mixtures by combining databases for each proteome from UniProtKB (44–46). The UPS1 and UPS2 FASTA database was downloaded from Sigma (47).

The mass spectrometry proteomics data have been deposited to the ProteomeXchange Consortium via the PRIDE (48) partner repository with the dataset identifier PXD030706 and 10.6019/PXD030706.

### 6.4 Software

The software was written in Python 3.8. The key libraries used were AlphaTims (49) for loading the raw data, Pandas 1.3.1 for data filtering and interface file input/output, scipy 1.6.1 and numpy 1.19.5 for signal processing, ms_deisotope 0.0.22 for spectra deconvolution, and Ray 1.5.2 for parallel processing. We used comet and percolator from crux 4.0 for searching the detected features against a FASTA peptide database.

We designed the software to implement distinct steps and to integrate with each other using file-based interfaces, usually Pandas dataframes serialised to the filesystem as feather files. The Feather format (https://arrow.apache.org/docs/python/feather.html) was used for its performance on large tables due to its columnar serialisation strategy, and for file portability between Python and R. Much of the bulk processing of samples to prove the algorithms was done on 48-core AWS EC2 memory- and compute-optimised instances running Ubuntu 20.04. As we aim to make this software accessible on commodity hardware, later validation work was performed on a PC with a 12-core Intel i7 6850K processor and 64 GB of memory running Ubuntu 20.04.

Readers are encouraged to browse the source code in the GitHub repository (DOI 10.5281/zenodo.6513126) for a detailed understanding of the algorithms and implementation approach. The Jupyter notebooks developed to generate the figures in this paper are also available in the repository.

## 7 Acknowledgments

The authors gratefully acknowledge the contributions of Sven Brehmer, Michael Krause, and Oliver Raether at Bruker for discussions and guidance about processing timsTOF data, and Jens Decker at Bruker for helpful feedback on a draft of the manuscript.

